# Comparing *in silico* and *in vitro* methods for classification of BCS II and CYP3A4 and MDR-1 substrate specificity

**DOI:** 10.1101/2022.12.13.520246

**Authors:** Urban Fagerholm

## Abstract

**Background:** Previous work has shown considerable laboratory variability of Biopharmaceutics Classification System (BCS) classification, efflux ratio in intestinal cell lines and cytochrome P450 (CYP450)-metabolism pathways. Such variability and inconsistency create uncertainty in predictions of human clinical pharmacokinetics and the pharmacokinetic optimization process and is a problem when developing corresponding *in silico* methods.

**Objectives and Methodology:** One objective of the study was to quantify the degree of laboratory inconsistency for BCS II-classing, MDR-1 and CYP3A4 substrate specificity (substrate/non-substrate). Another objective was to predict BCS II-classing, MDR-1 and CYP3A4 substrate specificity using *in silico* methodology and compare results to laboratory data/classifications.

**Results and Discussion:** 27 BCS II-classified drugs (with non-contradictory BCS-classing in various sources) were found. 17 (63 %) had an *in vivo* fraction absorbed (f_a_) of ≥90 % and belong to *in vivo* BCS I. With *in silico* methodology, 74 % correct BCS-classing was reached for the same set of compounds. The mean prediction error for f_a_ was 1.2-fold. MDR-1 and CYP3A4 substrate specificities were collected for 346 and 808 compounds, respectively. For MDR-1, 143 of the compounds had reported data in at least two studies, and out of these, 49 (34 %) and 18 (13 %) had contradictory (reported as both substate and non-substrate) and uncertain substrate specificities, respectively. For CYP3A4, 42 (9.8 %) out of 427 compounds showed inconsistency between laboratories. With *in silico* methodology, MDR-1 and CYP3A4 classification predictions were incorrect for 13 and 15 % of compounds.

**Conclusion:** The results show considerable variability/inconsistency for BCS II-classing (63 % inconsistency between BCS II-classing and *in vivo* f_a_) and MDR-1 (34 % inconsistency between sources) and CYP3A4 (10 % inconsistency between sources) substrate specificities. Corresponding estimates obtained with *in silico* methodology are 22, 13 and 15 %, respectively, demonstrating the power and applicability of such technology.

## INTRODUCTION

A previous investigation demonstrated considerable differences in solubility, permeability and efflux ratio (ER) in intestinal cell lines, Biopharmaceutics Classification System (BCS) classification and cytochrome P450 (CYP450)-metabolism pathways between and within laboratories (Fagerholm 2022). Such variability creates uncertainty in predictions of human clinical pharmacokinetics (PK) and in the PK-optimization process and is problematic for *in silico* method development.

MDR-1 (P-pg) and CYP3A4 are among the most important transporters and metabolic enzymes for drugs and drug candidates. For example, they contribute to lowering the oral bioavailability of many compounds, especially for compounds with limited passive permeability and high intrinsic metabolic clearance. A high variability/uncertainty of MDR-1 and CYP3A4 substrate specificity assays may therefore cause great uncertainty in PK-predictions of such compounds.

BCS-classification is used to in order to predict and plan clinical bioequivalence and food-drug interaction studies. Incorrect BCS-classification from *in vitro* data may lead to the planning and performance of unnecessary or unnecessarily large clinical studies or unexpected problems with bioinequivalence.

One objective of the study was to quantify the degree of laboratory inconsistency for BCS II-classing, MDR-1 and CYP3A4 substrate specificity (substrate/non-substrate). Another objective was to predict BCS II-classing, MDR-1 and CYP3A4 substrate specificity using *in silico* methodology and compare results to laboratory data/classifications.

## METHODOLOGY

### BCS II, MDR-1 and CYP3A4 data

BCS-classification and fraction absorbed (f_a_) data were taken from Pham-The et al. (2013). Only compounds with clear proposed BCS II were selected (compounds with different BCS-classing in various sources were excluded).

MDR-1 and CYP3A4 substrate specificity data (substrate/non-substrate) were taken from Bikadi et al. 2011 (MDR-1 substrates and non-substrates collected from the literature), Broccatelli et al. 2012 (a collection of ERs in MDCK-MDR1 cell lines), Sato et al. 2021 (ERs in MDCK-MDR1 cell line), Lin and Skolnik et al. 2010 and 2011 (ERs in MDCK-MDR1 and Caco-2 cell lines), Hayeshi et al. 2008 (ERs in Caco-2 cell line), Carbon-Mangels and Hutter 2011 (CYP3A4 substrate specificity), Yap and Chen 2005 (CYP3A4 substrate specificity) and Bioinformatics 2011 (CYP3A4 substrate specificity). In cases of missing limits for MDR-1, ER limits for substrate and non-substate were set to >1.3 and ≤1.

### *In silico* prediction methodology

The ANDROMEDA prediction software was used for prediction of f_a_ and BCS-class. For MDR-1 and CYP3A4 substrate specificity, quantitative structure–activity relationship*/*partial least square discriminant analysis (QSAR/PLS-DA) and quantitative structure–activity relationship*/*support vector machine (QSAR/SVM) models were applied (Fagerholm et al. 2021)

## RESULTS & DISCUSSION

### BCS II

27 BCS II-classified drugs (with non-contradictory BCS-classing in various sources) were found. 17 (63 %) of them had an *in vivo* f_a_ of ≥90 % and belong, therefore, to *in vivo* BCS I (Table 1; Figure 1).

**Table 1.**
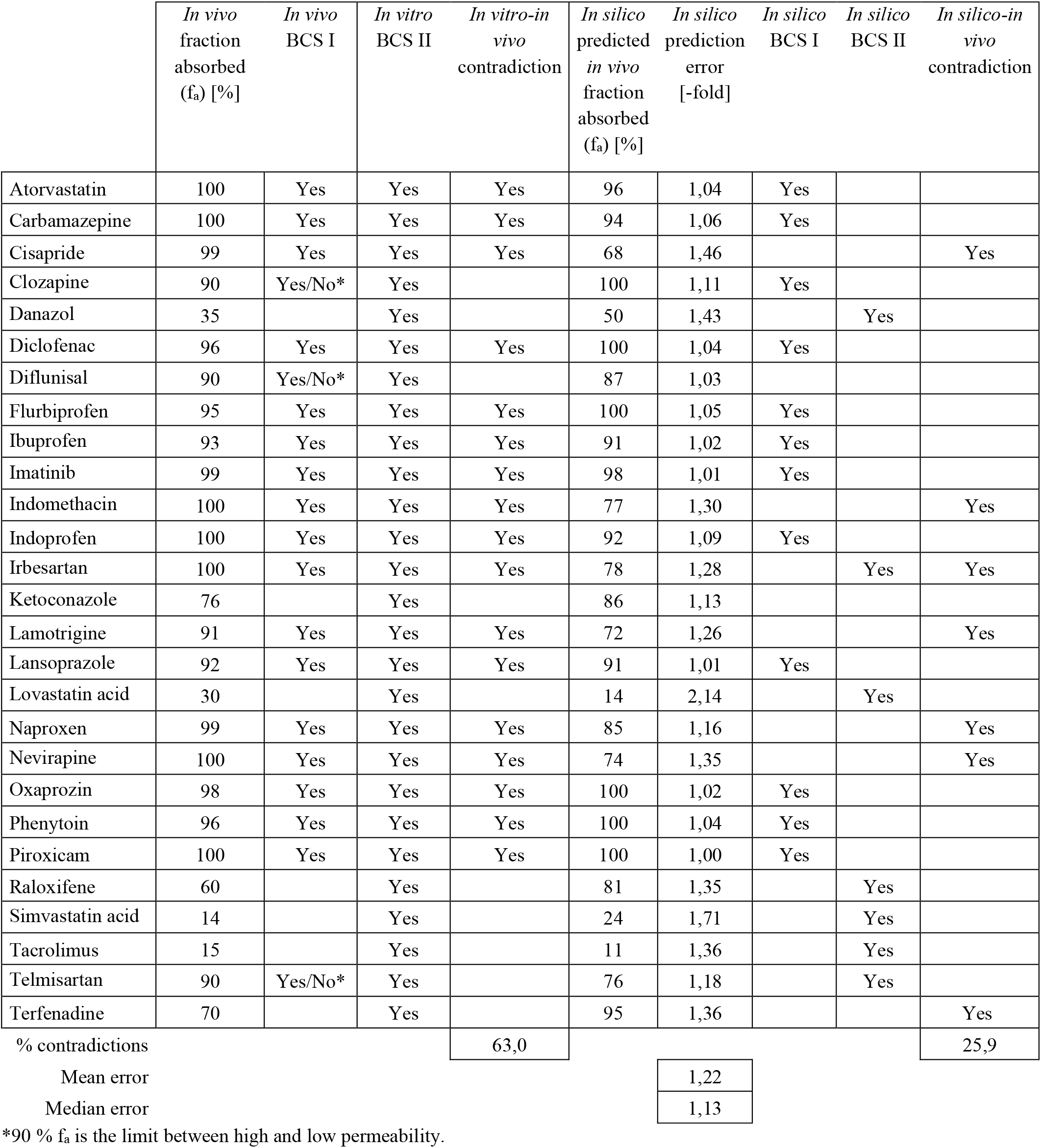
Results for BCS-classifications.

|  | *In vivo*<br>fraction<br>absorbed<br>(f<sub>a</sub>) [%] | *In vivo*<br>BCS I | *In vitro*<br>BCS II | *In vitro-in*<br>*vivo*<br>contradiction | *In silico*<br>predicted<br>*in vivo*<br>fraction<br>absorbed<br>(f<sub>a</sub>) [%] | *In silico*<br>prediction<br>error<br>[-fold] | *In silico*<br>BCS I | *In silico*<br>BCS II | *In silico-in*<br>*vivo*<br>contradiction |
| --- | --- | --- | --- | --- | --- | --- | --- | --- | --- |
| Atorvastatin | 100 | Yes | Yes | Yes | 96 | 1,04 | Yes |  |  |
| Carbamazepine | 100 | Yes | Yes | Yes | 94 | 1,06 | Yes |  |  |
| Cisapride | 99 | Yes | Yes | Yes | 68 | 1,46 |  |  | Yes |
| Clozapine | 90 | Yes/No* | Yes |  | 100 | 1,11 | Yes |  |  |
| Danazol | 35 |  | Yes |  | 50 | 1,43 |  | Yes |  |
| Diclofenac | 96 | Yes | Yes | Yes | 100 | 1,04 | Yes |  |  |
| Diflunisal | 90 | Yes/No* | Yes |  | 87 | 1,03 |  |  |  |
| Flurbiprofen | 95 | Yes | Yes | Yes | 100 | 1,05 | Yes |  |  |
| Ibuprofen | 93 | Yes | Yes | Yes | 91 | 1,02 | Yes |  |  |
| Imatinib | 99 | Yes | Yes | Yes | 98 | 1,01 | Yes |  |  |
| Indomethacin | 100 | Yes | Yes | Yes | 77 | 1,30 |  |  | Yes |
| Indoprofen | 100 | Yes | Yes | Yes | 92 | 1,09 | Yes |  |  |
| Irbesartan | 100 | Yes | Yes | Yes | 78 | 1,28 |  | Yes | Yes |
| Ketoconazole | 76 |  | Yes |  | 86 | 1,13 |  |  |  |
| Lamotrigine | 91 | Yes | Yes | Yes | 72 | 1,26 |  |  | Yes |
| Lansoprazole | 92 | Yes | Yes | Yes | 91 | 1,01 | Yes |  |  |
| Lovastatin acid | 30 |  | Yes |  | 14 | 2,14 |  | Yes |  |
| Naproxen | 99 | Yes | Yes | Yes | 85 | 1,16 |  |  | Yes |
| Nevirapine | 100 | Yes | Yes | Yes | 74 | 1,35 |  |  | Yes |
| Oxaprozin | 98 | Yes | Yes | Yes | 100 | 1,02 | Yes |  |  |
| Phenytoin | 96 | Yes | Yes | Yes | 100 | 1,04 | Yes |  |  |
| Piroxicam | 100 | Yes | Yes | Yes | 100 | 1,00 | Yes |  |  |
| Raloxifene | 60 |  | Yes |  | 81 | 1,35 |  | Yes |  |
| Simvastatin acid | 14 |  | Yes |  | 24 | 1,71 |  | Yes |  |
| Tacrolimus | 15 |  | Yes |  | 11 | 1,36 |  | Yes |  |
| Telmisartan | 90 | Yes/No* | Yes |  | 76 | 1,18 |  | Yes |  |
| Terfenadine | 70 |  | Yes |  | 95 | 1,36 |  |  | Yes |
| % contradictions |  |  |  | 63,0 |  |  |  |  | 25,9 |
| Mean error |  |  |  |  |  | 1,22 |  |  |  |
| Median error |  |  |  |  |  | 1,13 |  |  |  |

\*90 % f<sub>a</sub> is the limit between high and low permeability.

**Figure 1.**
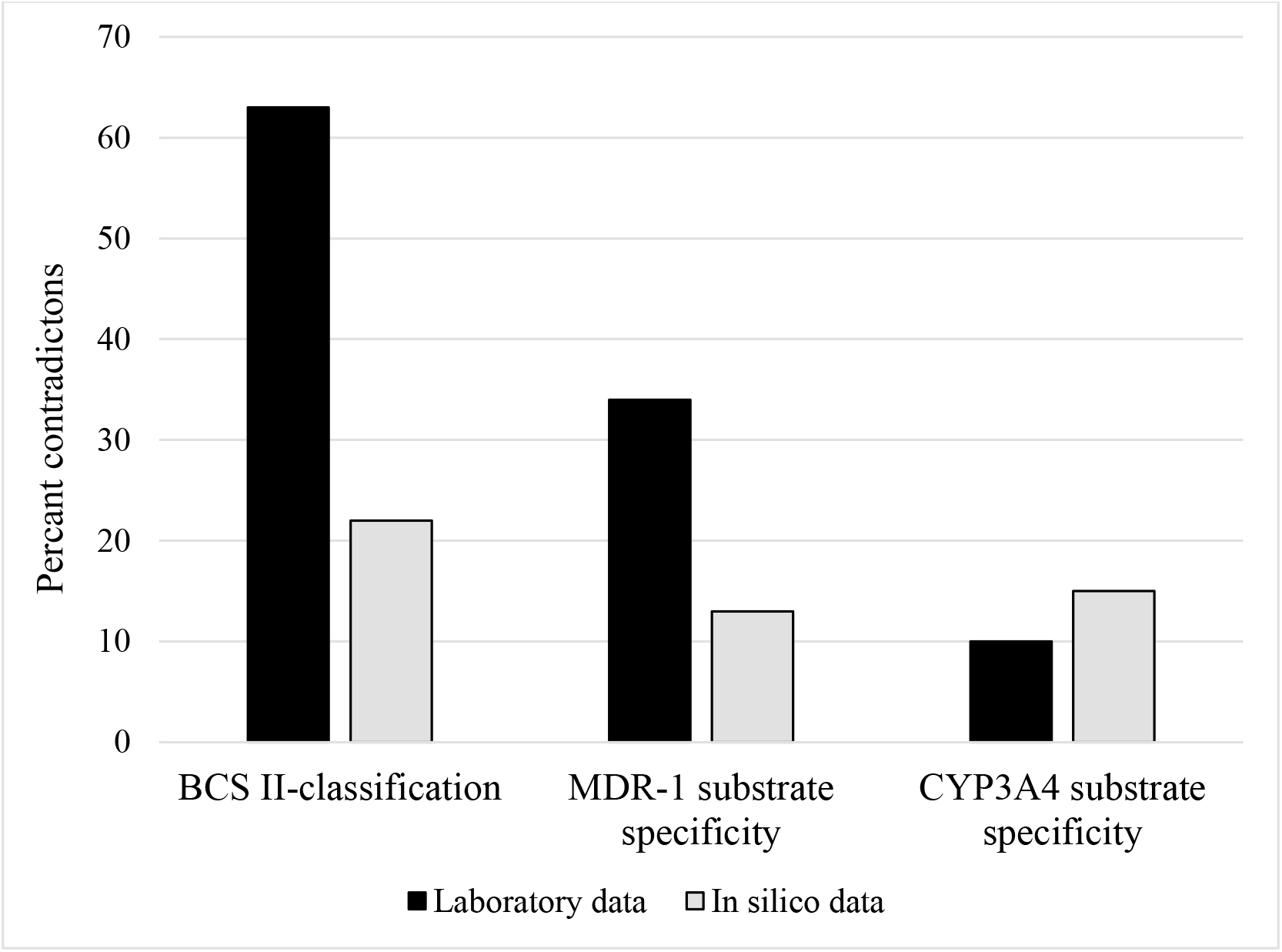
The percentages of contradictions for BCS II-, MDR-1- and CYP3A4-classifications with laboratory data (different data between laboratories/sources) and *in silico* predictions (difference between predicted and observed).

With *in silico* methodology, 74 % correct BCS-classing was reached for the same 27 compounds. The data set includes 7 compounds with predicted *in vivo* dissolution of less than 80 % and 7 compounds with a f_a_ of 85-95 % (±5 % from the limit between high and low permeability), which is challenging for predictions and BCS-classifications. The mean and median prediction errors for f_a_ were 1.22- and 1.13-fold, respectively.

### MDR-1 and CYP3A4 data

MDR-1 and CYP3A4 substrate specificities were collected for 346 and 808 compounds, respectively. For both MDR-1 and CYP3A4 about 60 % compounds were reported as substrates. For MDR-1, 143 of the compounds had reported data in at least two studies, and out of these, 49 (34 %) and 18 (13 %) had contradictory (reported as both substate and non-substrate) and uncertain substrate specificities (Figure 1). For CYP3A4, 42 (9.8 %) out of 427 compounds showed inconsistency between laboratories (Figure 1). Thus, the results show considerable interlaboratory variability/inconsistency for MDR-1 (53 % consistency) and CYP3A4 (90 % consistency) substrate specificities.

With *in silico* methodology, MDR-1 and CYP3A4 classification predictions were incorrect for 13 and 15 % of compounds (for each model data for several hundreds of compounds was used for the analysis).

The considerable variability/inconsistency between laboratories and experimental set-ups is likely to impact human clinical PK-predictions and *in silico* method development. One way to enhance the chance to correctly determine MDR-1 and CYP3A4 substrate specificities it to perform experiments using multiple methodologies and different conditions/set-ups. This is of particular importance for compounds with limited passive permeability (where MDR-1 efflux has greater impact) and low metabolic stability (where considerable gut-wall extraction may occur).

Based on these findings it is likely that similar results are expected for other important transporters and CYPs, such as BCRP, MRP2, CYP2C9 and CYP2D6.

In conclusion, the results show considerable variability/inconsistency for BCS II-classing (63 % inconsistency between BCS II-classing and *in vivo* f_a_) and MDR-1 (34 % inconsistency between sources) and CYP3A4 (10 % inconsistency between sources) substrate specificities. Corresponding estimates obtained with *in silico* methodology are 26, 13 and 15 %, respectively, which demonstrates the power and applicability of such technology.

## Notes

### Competing Interest Statement

Urban Fagerholm declares shares in Prosilico AB, a Swedish company that develops solutions for human clinical ADME/PK predictions.

